# Depletion of SMN Protein in Mesenchymal Progenitors Impairs the Development of Bone and Neuromuscular Junction in Spinal Muscular Atrophy

**DOI:** 10.1101/2023.10.06.561282

**Authors:** Sang-Hyeon Hann, Seon-Yong Kim, Ye Lynne Kim, Young-Woo Jo, Jong-Seol Kang, Hyerim Park, Se-Young Choi, Young-Yun Kong

**Affiliations:** School of Biological Sciences, Seoul National University, Seoul 08826, Republic of Korea; Department of Physiology, Dental Research Institute, Seoul National University School of Dentistry, Seoul, 03080, Republic of Korea

## Abstract

Spinal Muscular Atrophy (SMA) is a neuromuscular disorder characterized by the deficiency of the survival motor neuron (SMN) protein, which leads to motor neuron dysfunction and muscle atrophy. In addition to the requirement for SMN in motor neurons, recent studies suggest that SMN deficiency in peripheral tissues plays a key role in the pathogenesis of SMA. Using limb mesenchymal progenitor cells (MPCs)-specific SMN-depleted mouse models, we reveal that SMN reduction in chondrocytes and fibro-adipogenic progenitors (FAPs) derived from limb MPCs causes defects in the development of bone and neuromuscular junction (NMJ), respectively. We showed that impaired growth plate homeostasis, which causes skeletal growth defects in SMA, is due to reduced IGF signaling from chondrocytes rather than the liver. Furthermore, the reduction of SMN in FAPs resulted in abnormal NMJ maturation, altered release of neurotransmitters, and NMJ morphological defects. Transplantation of healthy FAPs rescued the morphological deterioration. Our findings highlight the significance of mesenchymal SMN in neuromusculoskeletal pathogenesis in SMA and provide insights into potential therapeutic strategies targeting mesenchymal cells for the treatment of SMA.

## Introduction

The survival motor neuron (SMN) protein is a crucial component of the spliceosome complex and is essential for the proper function of all cell types (Mercuri et al., 2022). Deficiency in SMN protein disrupts the formation of spliceosome complexes, ultimately causing splicing defects in multiple genes. Mutations in the *SMN1* gene, which encodes the SMN protein, give rise to the neuromuscular disorder Spinal Muscular Atrophy (SMA). SMA is characterized by neuromuscular junctions (NMJ) disruption, muscular atrophy, and alpha motor neuron loss (Mercuri et al., 2022; Burghes et al., 2009). The severity of the disease in humans correlates with the copy number of *SMN2*, which is a paralog of *SMN1* in humans. *SMN2* primarily produces less functional exon 7-deleted SMN protein and rarely generates a limited quantity of functional full-length SMN protein via alternative splicing. SMA patients are classified into types 0 through 4 based on the severity and timing of disease onset, predominantly determined by the number of copies of the *SMN2* gene they possess. More than 50% of SMA patients are categorized as type 1, characterized by muscle defects with proximal muscle atrophy during infancy, eventually resulting in death within a few years (Mercuri et al., 2022).

Previous studies suggest that the onset of SMA is mainly attributed to SMN loss in motor neurons (Monani et al., 2000; Burghes et al., 2009). Nevertheless, motor neuron-specific SMN deficiency in the SMA mouse model exhibits relatively mild phenotypes compared to whole-body SMA mouse models (Park et al., 2010; McGovern et al., 2015). Furthermore, restoring SMN to motor neurons in SMA mouse models result in only partial rescue in lifespan and neuromuscular defects (Passini et al., 2010; Martinez et al., 2012; McGovern et al., 2015; Besse et al., 2020). Systemic administration of antisense oligonucleotide (ASO), which corrects *SMN2* splicing to restore SMN expression, significantly prolongs survival compared to CNS administration (Hua et al., 2011). In the mouse treated with a systemically delivered ASO, blocking the effect of ASO in the CNS by a complementary decoy did not have any detrimental effect on survival, motor function, or NMJ integrity (Hua et al., 2015). These studies suggest that peripheral SMN plays a crucial part in SMA pathology. Overall, investigating the impacts of SMN depletion in peripheral tissues is critical for alleviating neuromuscular impairments and increasing life expectancy in SMA.

In SMA patients, bone growth retardation has been observed (De-Amicis et al., 2021; Kipoglu et al., 2022; Hensel et al., 2020). Studies using whole-body SMA mouse models have revealed that this is caused by diminished growth plate chondrocyte density and endochondral ossification defects, independent of muscle atrophy (Hensel et al., 2020). However, it is still unclear whether these defects result from the SMN ablation in chondrocytes, or from a decline in liver-derived IGF in SMA patients and mice (Hua et al., 2011; Yesbek-Kaymaz et al., 2016). In severe SMA mice, the serum levels of IGF decreased by approximately 60% or became undetected (Hua et al., 2011; Murdocca et al., 2012). The decreased serum IGF levels were attributed to decreased expression of liver genes, including *Igf1*, IGF binding, and ternary complex protein gene *Igfals* and *Igfbp3*. The previous studies suggest that the liver is the primary origin of systemic IGF, as demonstrated by the liver-specific deletion of *Igf1* and the knockout of *Igfals*, which is mostly expressed in the liver (Yakar et al., 1999; Yakar et al., 2002). The double KO mice exhibited a 90% decrease in serum IGF levels and displayed a phenotype of shortened femur length and growth plates. It is thus possible that the decrease in serum IGF levels, resulting from reduced liver IGF pathway genes in SMA mice, has also played a role in the observed bone growth defect.

Mesenchymal progenitor cells (MPCs) derived from the lateral plate mesoderm (LPM) differentiate into various types of limb mesenchymal cells, including bone, cartilage, and intramuscular mesenchymal cells like fibro-adipogenic progenitors (FAPs) (Nassari et al., 2017). Recent studies have revealed the role of FAPs in skeletal muscle homeostasis, as reducing the number of FAPs resulted in diminished muscle regeneration capacity, long-term muscle atrophy, and NMJ denervation (Wosczyna et al., 2019; Uezumi et al., 2021). Our latest research demonstrated that FAPs-specific deficiency of *Bap1*, one of the deubiquitinases, leads to NMJ defects (Kim et al., 2022). These recent findings raise the possibility that FAPs may have a specific role in the pathogenesis of neuromuscular diseases such as SMA. However, it has not been studied if depletion of SMN in FAPs can lead to SMA-like neuromuscular pathology.

In this study, we crossed a limb MPCs-specific Cre mouse with a floxed Smn exon7 mouse carrying multiple copies of the human *SMN2* gene, allowing us to examine the impact of mesenchymal SMN reduction on SMA pathogenesis. Our findings indicate that the SMN reduction in FAPs, similar to the extent of severe SMA, causes altered NMJ development like SMA. Additionally, in growth plate chondrocytes, it leads to skeletal growth abnormalities due to defects in autocrine and paracrine signaling of IGF.

## Results

### Bone growth restriction due to growth plate cell-autonomous defects caused by MPCs-specific SMN depletion

To investigate the effects of SMN reduction within MPCs in SMA pathogenesis, we crossed *Smn^f7/f7^* mice, which possess loxP sites flanking exon 7 of the *Smn* gene (Frugier et al., 2000), with *Prrx1^Cre^* mice. This produced Smn^ΔMPC^ mice (*Prrx1^Cre^; Smn^f7/f7^*) that lacked the *Smn* gene specifically in *Prrx1^Cre^*-expressed limb MPCs that give rise to bone, cartilage, and FAPs (Logan et al., 2002; Leinroth et al., 2022). To ascertain whether mutant mice carrying the *SMN2* gene, like SMA patients, present pathological phenotypes, we additionally generated SMN2 2-copy Smn^ΔMPC^ (*Prrx1^Cre^; Smn^f7/f7^; SMN2^+/+^)* and SMN2 1-copy Smn^ΔMPC^ (*Prrx1^Cre^; Smn^f7/f7^; SMN2^+/0^*) mice. Control littermates that lacked *Prrx1^Cre^* were used as controls for comparison.

SMN2 0-copy Smn^ΔMPC^ mice died within 24 hours after birth. Regions where limbs should have formed at E18.5 only had rudimentary limb structures (Figure 1—figure supplement 1A). Furthermore, since the upper head bone did not cover the brain, it was directly attached to the skin and protruded. These observations can be attributed to *Prrx1^Cre^*-mediated SMN deletion in the LPM-derived limb MPCs and the craniofacial mesenchyme, which is accountable for the formation of calvarial bone (Wilk et al., 2017). To investigate whether the lack of SMN proteins in MPCs is responsible for bone development abnormalities, we performed alcian blue and alizarin red staining on E18.5 SMN2 0-copy Smn^ΔMPC^ mutants to analyze the structure of bones and cartilage. The appendages displayed restricted bone and cartilage formations, with scarcely discernible femur and tibia (Figure 1—figure supplement 1B-C). In the cranial region, there was an absence of both cartilage and bone at the location of the calvarial bone, with the parietal bone entirely missing and partially absent frontal bone (Figure 1—figure supplement 1D). Additionally, the sternum, which is one of the bones originating from the LPM (Sheng et al., 2015), was shorter than the control (Figure 1—figure supplement 1E).

The SMN2 2-copy Smn^ΔMPC^ mice, carrying two homologous *SMN2* genes, did not show any discernible differences from the *Prrx1^Cre^*-negative control littermates into adulthood. However, the SMN2 1-copy Smn^ΔMPC^ mice exhibited reduced body size and shorter limb length compared to the SMN2 1-copy control (*Smn^f7/f7^; SMN2^+/0^*). To assess postnatal bone growth defects observed in SMA patients and mouse models, we conducted micro-CT analysis on femurs obtained from postnatal day 14 (P14) SMN2 1-copy Smn^ΔMPC^ and SMN2 2-copy Smn^ΔMPC^ mice (Figure 1A). The 3D reconstruction image showed that the SMN2 1-copy mutant femur was smaller than the WT control and SMN2 2-copy mutant, and secondary ossification center is denied. The longitudinal virtual section view displayed reduced trabecular bone in the SMN2 1-copy mutant femur. CT analysis data showed that SMN2 1-copy mutants exhibited reduced femur diaphysis length, diameter, and trabecular bone volume compared to the control group, indicating growth plate-dependent endochondral ossification defects (Figure 1B-D). We then examined femoral bone thickness and diaphyseal bone mineral density (BMD) to determine whether mineralization was normal after bone formation. The thickness of the bone in the mutants did not differ significantly from the control, suggesting that bone mineralization was intact (Figure 1E and Figure 1—figure supplement 2A). Unexpectedly, BMD slightly increased in SMN2 1-copy mutants compared to the control group (Figure 1—figure supplement 2B). To assess the impact of osteoclasts and osteoblasts on diaphysis cortical bone mineralization, we utilized *Itgb3* immunofluorescence as the osteoclast marker and toluidine blue staining for imaging bone-attached osteoblasts (Romeo et al., 2019; Colaianni et al., 2015). Osteoclast and osteoblast density did not significantly differ between the SMN2 1-copy mutant and the control (Figure 1—figure supplement 2C-E). The higher BMD may be attributed to greater mechanical stress caused by the shorter femur supporting the weight of the body, consistent with prior research indicating that elevated mechanical force leads to higher BMD in the femur (Hoxha et al., 2014; Ike et al., 2015). Nevertheless, the decrease in bone growth without apparent deterioration in bone mineralization of the femur of SMN2 1-copy Smn^ΔMPC^ mutants is consistent with findings from the whole-body SMA mouse model (Hensel et al., 2020). Collectively, these results suggest that mice carrying low copies of *SMN2*, with the SMN gene specifically deleted in MPCs, exhibit bone growth abnormalities, while osteoblast and osteoclast populations remain unaffected.

**Figure 1.**
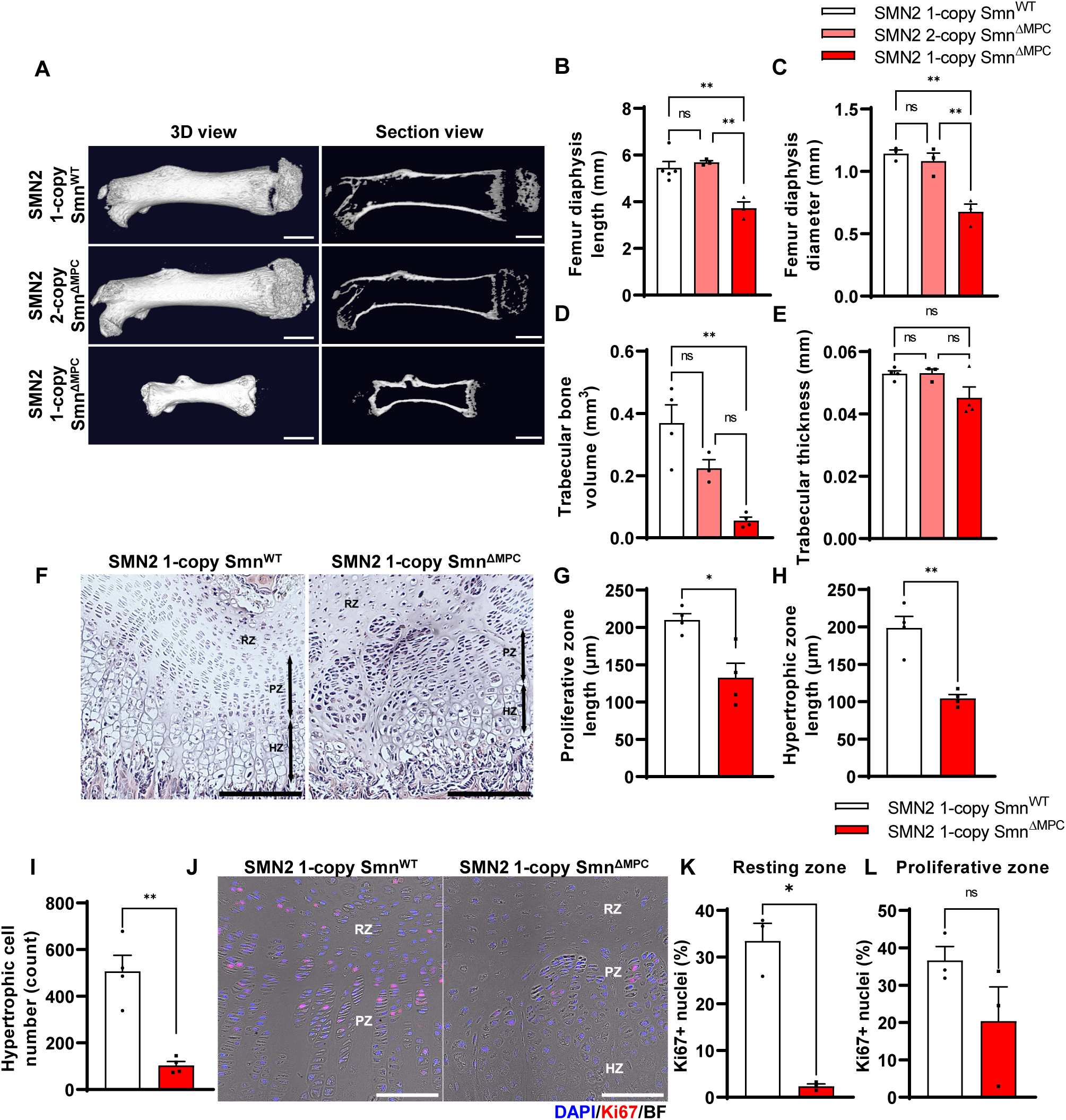
Skeletal growth abnormalities and altered growth plate homeostasis in SMN2 1-copy Smn^ΔMPC^ mice. (**A**) Representative 3D images and longitudinal section view of the ossified femur bone. Scale bars, 1 mm. (**B-C**) SMN2 1-copy mutant’s femurs showed reduced growth in diaphysis length and diameter, and (**D**) decreased trabecular bone volume. (**E**) Trabecular bone thicknesses were not significantly different between the control and mutant groups. The micro CT analysis was performed in femur diaphysis and metaphysis from SMN2 1-copy Smn^ΔWT^, SMN2 2-copy, and 1-copy Smn^ΔMPC^ mice at P14. 1-way ANOVA with Tukey’s post hoc test, n = 3-5 mice in each genotype (**B-E**). (**F)** Representative images of H&E staining in the distal femur growth plate of control and mutant mice with 1 copy of *SMN2* at P14. Scale bars, 100 μm. Resting zone (RZ), Hypertrophic zone (HZ) and proliferative zone (PZ). (**G-I**) Indicated by black arrows, the HZ and PZ lengths were reduced in SMN2 1-copy Smn^ΔMPC^ mice, and the hypertrophic cell number in a section of the 1-copy mutant was decreased (n = 4 mice in each genotype; Unpaired t-test with Welch’s correction). (**J)** Representative images of Ki67 immunostaining in the distal femur growth plate of control and mutant mice with 1 copy of *SMN2* at P14. Scale bars, 100 μm. (**K**) and decreased Ki67+ percentage in resting zone chondrocytes. (**L**) Ki67+ percentage in the proliferative zone was not significantly different between the control and mutant groups. n = 3 mice in each genotype; Unpaired t-test with Welch’s correction (**K-L**). ns; not significantly different. \**p* < 0.05; \*\**p* < 0.01. Error bars show s.e.m.

Various mesenchymal cells originating from the lateral plate mesoderm were known to contribute to bone formation, such as growth plate chondrocytes, osteoblasts, and bone marrow stromal cells (BMSCs). In SMN2 1-copy Smn^ΔMPC^ mutants, SMN may be deleted in all of these cells, suggesting that they all play a role in the bone growth abnormalities observed in 2 weeks mice. However, the population of Lepr+Cxcl12+ BMSCs, which makes up the majority of bone marrow stromal cells and is positive for Prrx1, was reported to be capable of forming new bone only in adults that are over 8 weeks old (Zhou et al., 2014; Matsushita et al., 2020). Consequently, it appears improbable that the observed phenotype at 2 weeks was due to SMN deletion in BMSCs. Furthermore, previous researchers revealed that primary osteoblasts from a severe SMA mouse model did not display notable differences from controls in an in vitro ossification test. And they did not observe any differences in bone voxel density and bone thickness in femurs at P3 severe SMA mice. This is supported by the absence of any bone thickness or BMD defects in the SMN2 1-copy mutant (Figure 1E and Figure 1-figure supplement 2A-B), and the unimpaired osteoblast population (Figure 1-figure supplement 2E). Thus, we conclude that the bone growth abnormalities observed in the 2-week-old SMN2 1-copy Smn^ΔMPC^ mutant are due to impaired endochondral ossification of growth plate chondrocytes.

To determine whether bone growth defects in SMN2 1-copy Smn^ΔMPC^ mutants arise from disrupted chondrocyte homeostasis at growth plates, we stained the femur distal growth plate of P14 mice with hematoxylin and eosin (Figure 1F). In line with earlier findings in whole-body SMA mice (Hensel et al., 2020), SMN2 1-copy Smn^ΔMPC^ mice exhibited shorter proliferative and hypertrophic zones compared to control mice (Figure 1G and H). Additionally, there was a significant reduction in the number of chondrocytes in the hypertrophic zone (Figure 1I). To investigate the decrease in chondrocyte proliferation and subsequent reduction in the proliferative and hypertrophic zone, we stained the proliferation marker Ki67 in the growth plate of both SMN2 1-copy control and mutant samples (Figure 1J). We then quantified the percentage of Ki67+ nuclei in the resting and proliferative zones (Figure 1K and L). Although there was no significant difference observed in the Ki67+ percentage of proliferative zone chondrocytes in the proceeding proliferation state, there was an absolute reduction in resting zone chondrocyte proliferation. The decreased proliferation rate in the resting zone could have impeded the transition to the proliferative zone. Our data indicate that adequate expression of SMN is essential for the homeostasis of chondrocytes at growth plates.

### Disruption of chondrocyte-derived IGF signaling in SMN2 1-copy Smn**^Δ^**^MPC^ mutants

The proliferation and differentiation of growth plate chondrocytes are regulated by systemic IGF (Shim et al., 2015; Karimian et al., 2012; Shantanam et al., 2018; Racine et al., 2020). Previous research suggested that a key factor contributing to the pathological phenotype in SMA is the lowered expression of the *Igf1/Igfbp3/Igfals* genes, which produce IGF and IGF-carrying proteins, in the liver. (Hua et al., 2011; Murdocca et al., 2012). As the IGF pathway proteins are downregulated in whole-body SMA mice, the bone growth defects observed in the mice have sparked debate as it remains unclear whether they are due to cell-autonomous defects by chondrocyte SMN reduction, or the low liver-derived IGF level (Hua et al., 2011; Tsai et al., 2014; Deguise et al., 2021; Hensel et al., 2015; Hensel et al., 2020).

To clarify this issue, we used Smn^ΔMPC^ mutant mice, which enabled us to investigate the effect of SMN depletion in chondrocytes on bone growth, while circumventing the impact of the endocrine signal by *Prrx1*-negative organ. To investigate the impact of IGF signaling on growth plate chondrocytes, we employed immunofluorescence to evaluate the percentage of p-AKT positive cells activated by the IGF-PI3K-AKT pathway in both SMN2 1-copy control and mutant femur distal growth plate (Figure 2A). Intriguingly, the percentage of p-AKT+ cells was significantly decreased in resting zone chondrocytes, but not in the proliferative zone, which aligns with the Ki67+ percentage (Figure 2B-C). Expectedly, the liver’s mRNA expression of IGF pathway genes, reported to be decreased in SMA mouse models and patients (Hua et al., 2011; Murdocca et al., 2012; Deguise., 2021; Sahashi et al., 2013), showed no difference when comparing controls to SMN2 1-copy Smn^ΔMPC^ mutants (Figure 2D). These findings indicate that the growth plate’s proliferation and hypertrophy in SMN2 1-copy mutants are affected by impairments in another AKT upstream signal rather than by liver-secreted systemic IGF.

**Figure 2.**
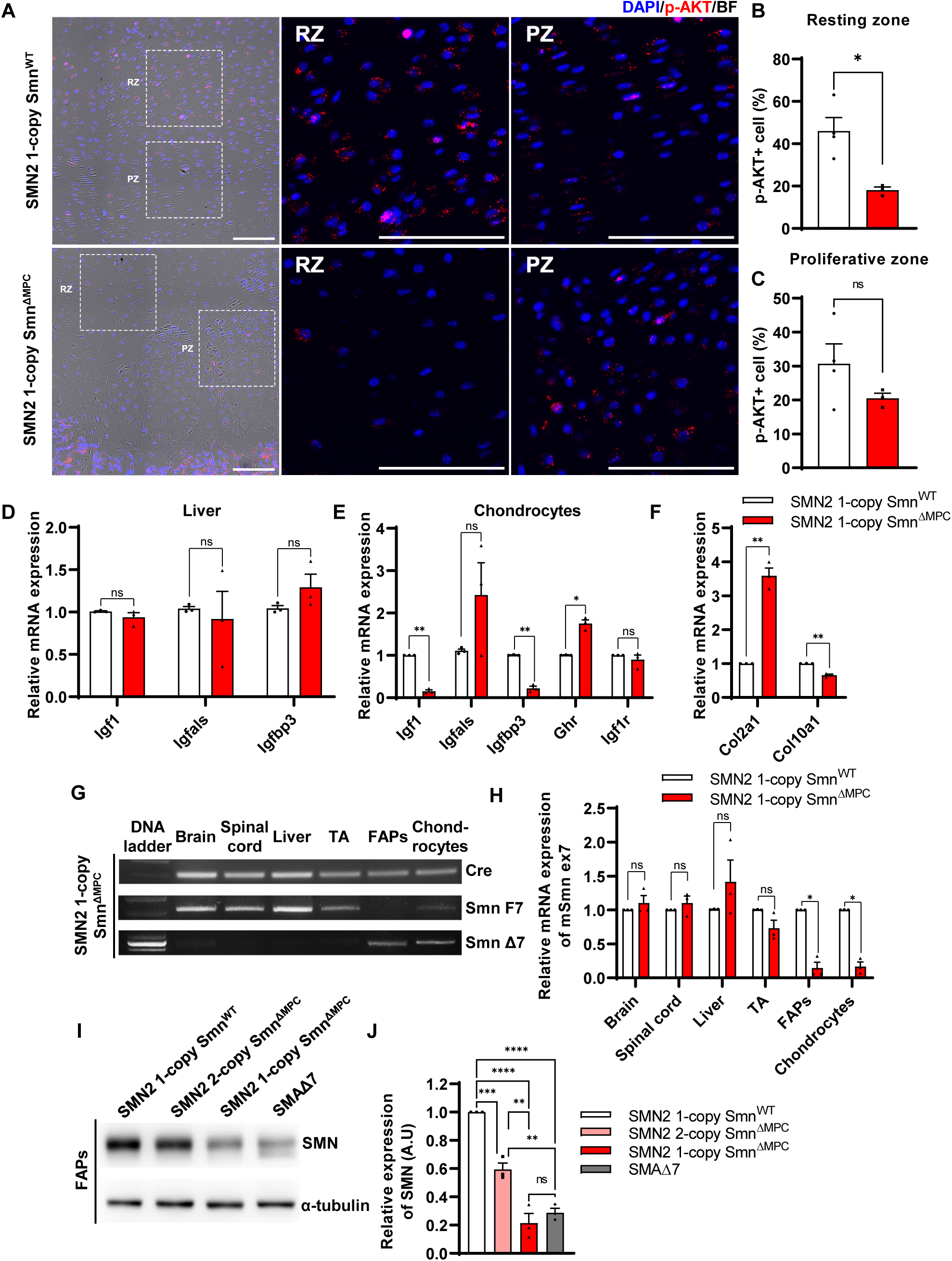
Decreased chondrocytes-derived IGF-AKT axis by limb mesenchymal cell-specific SMN depletion in SMN2 1-copy Smn^ΔMPC^ mice. (**A**) Representative images of p-AKT immunostaining in distal femur growth plate from mice at P14. Scale bars, 100 μm. (**B-C**) The p-AKT-positive percentage was decreased in the resting zone chondrocytes, not in the proliferative zone. (n = 3-4 mice in each genotype; Unpaired t-test with Welch’s correction) (**D**) Relative IGF axis mRNA expression in the livers of SMN2 1-copy control and mutant mice. The IGF pathway genes showed no difference when comparing controls to SMN2 1-copy Smn^ΔMPC^ mutants. (**E-F**) Relative IGF axis and chondrocyte differentiation marker mRNA expression in the chondrocytes of SMN2 1-copy control and mutant mice. The *Igf1, Igfbp3*, and hypertrophic marker *Col10a1* expression were decreased in SMN2 1-copy Smn^ΔMPC^ mutants. n = 3 mice in each genotype; Unpaired t-test with Welch’s correction (**D-F**). (**G**) Representative images of genomic PCR analysis from SMN2 1-copy Smn^ΔMPC^ mice tissues at P21. (**H**) qRT PCR analysis from tissues of SMN2 1-copy Smn^WT^ and Smn^ΔMPC^ mice at P21 (n = 3 mice in each genotype; Unpaired t-test with Welch’s correction). Deletion of Smn exon 7 was detected only in limb mesenchymal cells using genomic PCR (**G**) and full-length SMN mRNA expression (**H**). (**I**) Representative images of western blot analysis in cultured FAPs. SMN protein in FAPs of SMN2 1-copy Smn^ΔMPC^ mice exhibited a decrease comparable to that observed in the SMAΔ7 mice. (**J**) Relative SMN levels in cultured FAPs of the controls and mutants (n = 3 mice in each genotype; 1-way ANOVA with Tukey’s post hoc test). ns; not significantly different. \**p* < 0.05; \*\**p* < 0.01; \*\*\**p* < 0.001; \*\*\*\**p* < 0.0001. Error bars show s.e.m.

There are reports indicating that local IGF expression plays a crucial role in bone development through the growth plate, in addition to circulating IGF (Hallett et al., 2019; Racine et al., 2020). While most *Igf1*-null mice died before birth and had smaller tibial lengths than normal, liver-specific *Igf1*-deleted mice did not experience significant changes in body length or tibial length during postnatal growth, despite a 75% reduction in serum IGF-1 levels (Baker et al., 1993; Yakar et al., 1999). This indicates that local IGF in the growth plate is crucial for endochondral ossification, in addition to serum IGF. The chondrocyte-specific Igf1 knockout mouse demonstrated a reduction in postnatal body and femur length, and the chondrocyte-specific *Igf1r* knockout mouse demonstrated a significant reduction in bone growth, as well as a decrease in both growth plate proliferative and hypertrophic zone (Govoni et al., 2007; Wang et al 2011). A recent study revealed that resting zone chondrocytes in the growth plate serve as a major source of local IGF and activate the p-AKT pathway via autocrine and paracrine IGF signaling (Oichi et al., 2023). The study further revealed that cells that constitute bone and bone marrow, apart from chondrocytes, do not express *Igf1*, making chondrocytes the solitary source of local IGF. These suggest that the growth plate defects and the reduction of resting zone AKT phosphorylation in SMN2 1-copy mutants may be due to chondrocyte-secreted IGF deficiency. To confirm this hypothesis, we evaluated the expression of IGF-related genes in chondrocytes from both SMN2 1-copy control and mutant femur (Figure 2E). Indeed, *Igf1* and *Igfbp3* were greatly depleted in SMN2 1-copy mutant chondrocytes. It is hypothesized that the increased presence of *Ghr* may be in response to the reduction of *Igf1*. As IGF directly causes chondrocyte hypertrophy (Wang et al., 1999), we assess the mRNA expression of chondrocyte hypertrophic marker *Col10a1* and undifferentiated chondrocyte marker *Col2a1* (Figure 2F). The results show that *Col10a1* is decreased, while *Col2a1* is increased in the mutant. In Figure 1H-I, the hypertrophic cell reduction may be caused by a low local IGF level. Therefore, depletion of local IGF in the growth plate of SMN2 1-copy mutants may hinder chondrocyte progression to proliferation and hypertrophy, leading to aberrations in bone development. In Figures 1L and 2C, it is possible that the serum IGF could affect the remnant proliferative and p-AKT-positive cells in the growth plate via the vascularization of bone marrow. Based on these findings, we concluded that the bone growth defects observed in the whole-body SMA mouse model are mainly due to a depletion of localized IGF secretion by SMN-deleted chondrocytes.

### Mesenchymal cell-specific SMN reduction similar to severe SMA mouse model in SMN2 1-copy Smn**^Δ^**^MPC^ mutants

To confirm the specific deletion of *Smn* in limb mesenchymal cells, including chondrocytes and FAPs, we performed qRT-PCR for full-length SMN mRNA expressed by undeleted *Smn* and the *SMN2* alternative splicing in each tissue. Our findings indicate that full-length SMN mRNA expression in the brain, liver, skeletal muscle, and spinal cord of SMN2 1-copy Smn^ΔMPC^ mice at postnatal day 21 (P21) was similar to that of control mice, while it was significantly reduced in isolated FAPs and chondrocytes (Figure 2H). Additionally, we confirmed the presence of the SmnΔ7 variant, which is the exon 7-deleted form of *Smn* created by cre-lox mediated recombination, in both FAPs and chondrocytes by conducting genomic PCR on the SMN2 1-copy Smn^ΔMPC^ mutant (Figure 2G). Since SMN was not down-regulated in the tissues other than limb mesenchymal cells, non-mesenchymal cells were ruled out from being responsible for the phenotype observed in Smn^ΔMPC^ mutants.

Among our Smn^ΔMPC^ models, mice with two copies of *SMN2* exhibit similar bone development parameters as control mice without SMN deletion. However, mice with one copy of *SMN2* display bone pathological defects akin to SMA mouse models. This may be because the quantity of human full-length SMN protein produced by SMN2 2-copy, was sufficient to sustain SMN complex function in the SMN2 2-copy Smn^ΔMPC^ mutants, despite the absence of functional mouse SMN protein in limb mesenchymal cells. On the contrary, due to the lower expression of full-length SMN compared to SMN2 2-copy mutants, the SMN complex may not function properly in SMN2 1-copy mutant cells. We confirmed this by comparing the amount of full-length SMN protein in isolated FAPs from the hindlimbs of control, Smn^ΔMPC^ mutants, and a severe SMA mouse model (*Smn^-/-^; SMN2^+/+^; SMN*Δ*7^+/+^*; -SMAΔ7 mutants -) (Figure 2I). The data show that SMN2 1-copy Smn^ΔMPC^ mutants exhibited ∼80% reduction in SMN levels compared to the control group. The level of SMN protein in SMN2 1-copy Smn^ΔMPC^ mutants was similar to that in SMAΔ7 mutants. Conversely, SMN2 2-copy mutants display a decrease of approximately 40% in SMN protein levels compared to the control (Figure 2J). The moderately reduced expression of SMN is adequate to support regular bone development in SMN2 2-copy mutants. Taken together, our findings indicate that the reduction of mesenchymal SMN to levels comparable to that of the severe SMA mouse model causes SMA-like bone pathology in the SMN2 1-copy mutant.

### Abnormal NMJ maturation in SMN2 1-copy Smn**^Δ^**^MPC^ mutants

To investigate whether disabling SMN in FAPs results in SMA-like neuromuscular impairments, we assessed if NMJ phenotypes observed in SMA mouse models also occur in SMN2 1-copy Smn^ΔMPC^ mutants. Both severe and mild SMA mouse models exhibit impaired NMJ maturation markers, including plaque-like morphology of acetylcholine receptor (AChR) clusters, neurofilament (NF) varicosities, and poor terminal arborization (Kong et al., 2009; Martinez, 2012; Monani et al., 2003; Kariya et al., 2008). To evaluate the impact of SMN deficiency in FAPs on NMJ maturation, we evaluated the NMJ maturation markers in the TA muscles of control and SMN2 1-copy Smn^ΔMPC^ mutant mice at P21, a time when NMJ maturation is in progress (Figure 3A). Our examination revealed the presence of NF varicosities in SMN2 1-copy mutants as compared with control mice (Figure 3A and 3D). Additionally, the number of nerve branches was decreased and half of the total NMJs were poorly arborized in SMN2 1-copy mutants (Figure 3B-C). These presynaptic alterations are specific phenotypes in neurogenic atrophy like SMA. Unlike neurogenic atrophy, physiologic atrophy shows no differences in presynaptic morphology, such as nerve branching (Deschenes et al., 2006). This suggests that the NMJ phenotypes observed in SMN2 1-copy Smn^ΔMPC^ mutant mice are not caused by decreased muscle size and activity resulting from bone growth abnormalities. The morphology of AChR clusters shows that the mutants have more immature plaque-like NMJs than the controls’ pretzel-like structure (Figure 3E). Therefore, these findings indicate that SMN2 1-copy Smn^ΔMPC^ mutants exhibit NMJ maturation abnormalities common in SMA mouse models.

**Figure 3.**
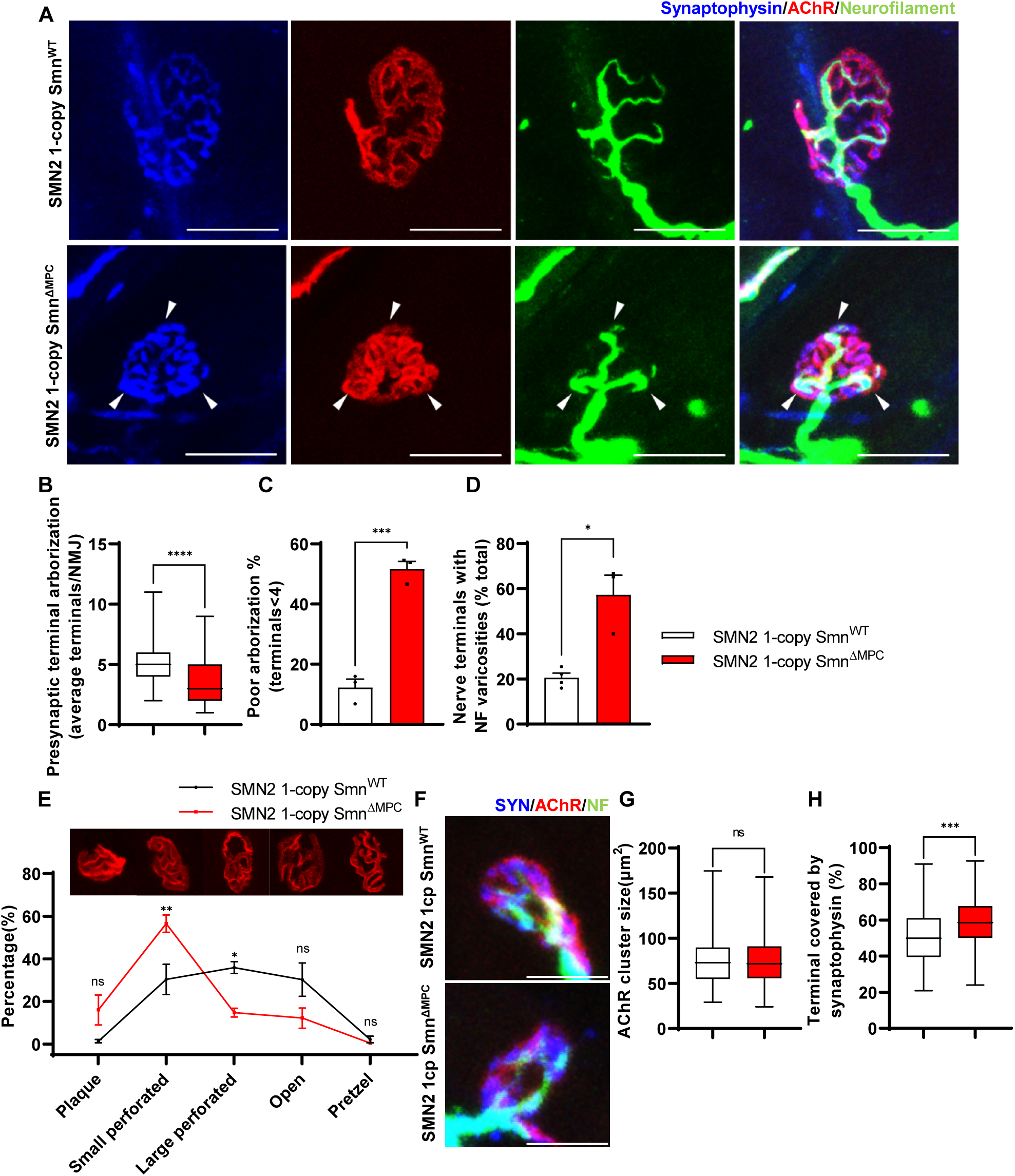
Aberrant postnatal NMJ maturation in SMN2 1-copy Smn^ΔMPC^ mice. (**A**) Immunostaining of NMJs in TA muscle of SMN2 1-copy Smn^WT^ and Smn^ΔMPC^ mice at P21 with anti-NF (Green), anti-synaptophysin (Blue), α-Btx staining AChR (Red). Scale bars, 20 μm. The confocal images of NMJs showed decreased presynaptic terminal branching and the existence of nerve terminal varicosities that were enlarged with NF (indicated by arrowheads) in the mutant. (**B**) The NMJs of the SMN2 1-copy mutant exhibited a significant decrease in presynaptic terminal arborization and (**C**) an increased percentage of poorly arborized NMJs (n = 3 mice in each genotype; Unpaired t-test with Welch’s correction). (**D**) The percentage of NMJs exhibiting NF varicosities was higher in the SMN2 1-copy mutant group than in the control group (n = 3-4 mice in each genotype; Unpaired t-test with Welch’s correction). (**E**) For quantification of the NMJ maturation stage, we classified NMJs into five distinct developmental stages (Plaque: Plaque-shaped endplate without any perforation; Small perforated: Plaque-shaped endplate with small perforations; Large perforated: Plaque-shaped endplate with large perforations; Open: C-shaped endplate; Pretzel: Pretzel-like shaped endplate) and then compared the frequency patterns of SMN2 1-copy control and mutant mice (n = 3 mice in each genotype; 2-way ANOVA with Tukey’s post hoc test). The NMJs of SMN2 1-copy mutants displayed plaque-like shapes, indicating that they were in the immature stage. (**F**) Immunostaining of NMJs in TA muscle of SMN2 1-copy Smn^WT^ and Smn^ΔMPC^ mice at P3 with anti-NF (Green), anti-synaptophysin (Blue), α-Btx staining AChR (Red). Scale bars, 10 μm. (**G**) There were no significant differences in AChR cluster size between the SMN2 1-copy control and mutant at P3 (n = 3-4 mice in each genotype; Unpaired t-test with Welch’s correction). (**H**) The ratio of the Synaptophysin area to the AChR area in NMJ was slightly higher in the SMN2 1-copy mutant at P3 (n = 3-4 mice in each genotype; Unpaired t-test with Welch’s correction). ns; not significantly different. \**p* < 0.05; \*\**p* < 0.01; \*\*\**p* < 0.001; \*\*\*\**p* < 0.0001. All box-and-whisker plots show the median, interquartile range, minimum, and maximum. For the box-and-whisker plots, range bars show minimum and maximum (**B, G, H**). For the bar and line graph, error bars show s.e.m(**C, D, E**).

### Undisturbed NMJ formation in neonatal SMN2 1-copy Smn**^Δ^**^MPC^ mutants

To determine whether any NMJ defects were present prior to juvenile NMJ maturation in SMN2 1-copy Smn^ΔMPC^, we examined NMJ formation in SMN2 1-copy Smn^ΔMPC^ mice at the neonatal stage on postnatal day 3. We evaluated the AChR and nerve terminal areas to assess postsynaptic and presynaptic development, respectively (Figure 3F). Measurements of AChR cluster size indicated no differences between control and SMN2 1-copy Smn^ΔMPC^ mice (Figure 3G). However, the area of AChR covered by nerve terminals was slightly larger in SMN2 1-copy Smn^ΔMPC^ (Figure 3H). We have no reasonable explanation for why the coverage is higher in the mutant. However, there does not appear to be abnormal development of the NMJ in the mutant, at least until the neonatal period. Therefore, we reasoned that SMN2 1-copy Smn^ΔMPC^ mutants began to exhibit deterioration in the NMJ maturation during the juvenile stage, following the intact neonatal development of NMJ.

### Aberrant NMJ morphology in the adult SMN2 1-copy Smn**^Δ^**^MPC^ mice

To evaluate the organization of NMJ after the conclusion of postnatal NMJ development, considering mesenchymal SMN expression, we examined NMJ morphology in the TA muscle of control, SMN2 1-copy, and 2-copy Smn^ΔMPC^ mice at postnatal day 56 (P56). Our analysis revealed that presynapses were fragmented in SMN2 1-copy mutants, resulting in a bouton-like morphology, in contrast to the control and SMN2 2-copy Smn^ΔMPC^ mice (Figure 4A). In SMN2 1-copy Smn^ΔMPC^ mice, a two-fold presynaptic fragmentation compared to the control was quantified, demonstrating nerve terminal shrinkage (Figure 4B). Additionally, NF ends displayed more severe varicosities than at P21 and were only connected to the proximal nerve by very thin NF, unlike control and SMN2 2-copy mice (Figure 4C). Remarkably, numerous presynaptic islands formed in SMN2 1-copy Smn^ΔMPC^ mice through the merging of fragmented presynapses and NF varicosity. In SMN2 1-copy mutants, AChR clusters displayed fragmented grape-shaped morphology that overlapped with nerve terminals, whereas control and SMN2 2-copy mice displayed pretzel-like structures (Figure 4D). These results suggest that defects in adult NMJ morphology occur when mesenchymal SMN protein is reduced to the extent of the SMN2 1-copy Smn^ΔMPC^ mutants.

**Figure 4.**
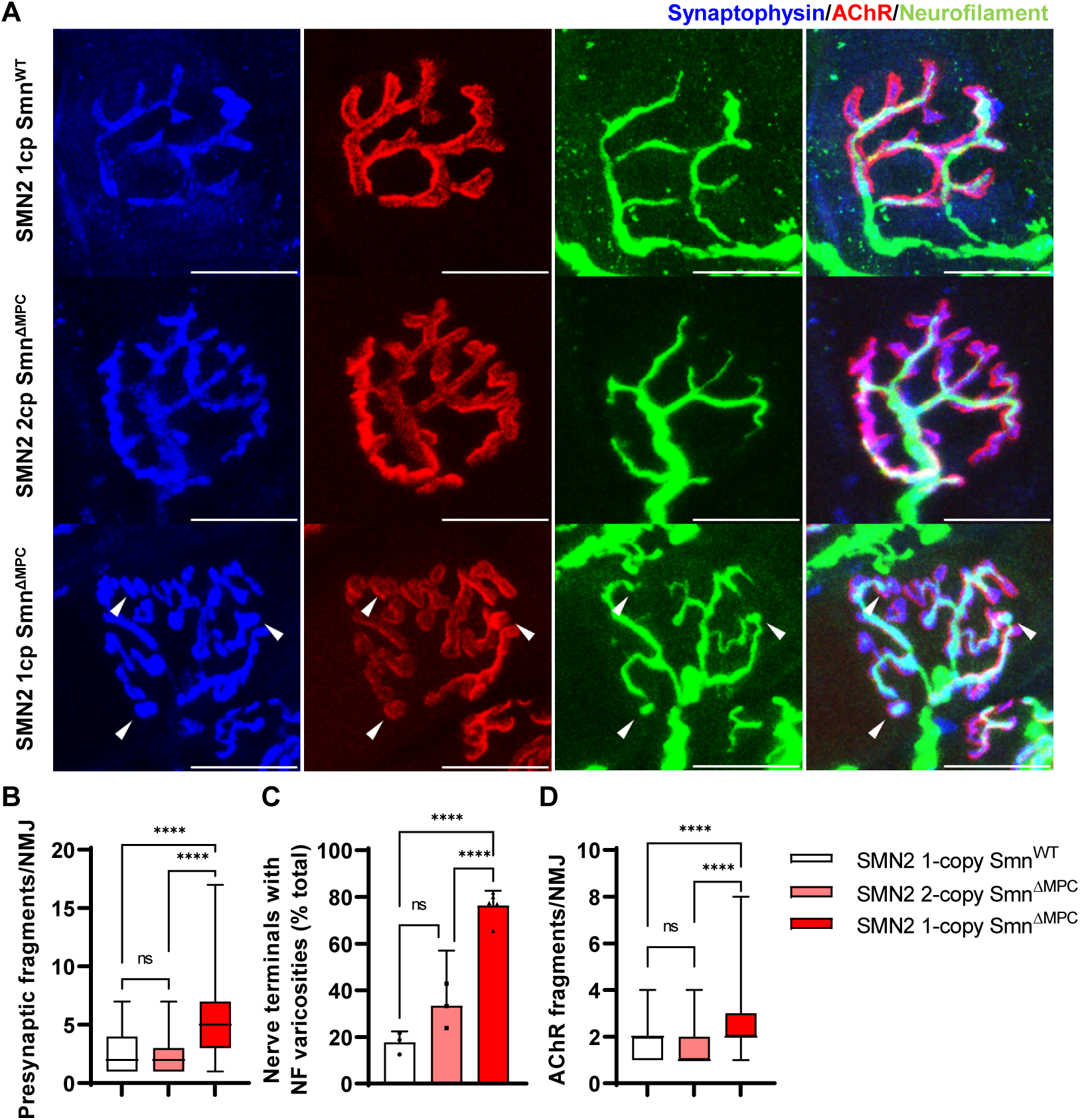
Morphological deterioration in NMJs of adult SMN2 1-copy Smn^ΔMPC^ mice. (**A**) Immunostaining of NMJs in TA muscle of SMN2 1-copy Smn^WT^, SMN 2-copy, and SMN2 1-copy Smn^ΔMPC^ mice at P56 with anti-NF (Green), anti-synaptophysin (Blue), α-Btx staining AChR (Red). Scale bars, 20 μm. The confocal images of NMJs showed fragmentation and bouton-like NF varicosities (indicated by arrowheads) in the SMN2 1-copy Smn^ΔMPC^ mice. (**B**-**D**) The NMJs of the SMN2 1-copy mutant displayed fragmented presynapse (**B**), endplate (**D**), and NF varicosities (**C**) compared to SMN2 1-copy Smn^WT^ and SMN2 2-copy Smn^ΔMPC^ mice (n = 3-5 mice in each genotype; Presynaptic fragments & AChR fragments: Brown– Forsythe and Welch ANOVA with Games-Howell’s test; NF varicosities: 1-way ANOVA with Tukey’s post hoc test). ns; not significantly different. \*\*\*\**p* < 0.0001. All box-and-whisker plots show the median, interquartile range, minimum, and maximum. For the box-and-whisker plots, range bars show minimum and maximum (**B, D**). For the bar graph, error bars show s.e.m. (**C**)

### Presynaptic neurotransmission alteration in SMN2 1-copy Smn**^Δ^**^MPC^ mutants

To investigate whether the morphologically aberrant NMJs of SMN2 1-copy Smn^ΔMPC^ mice have functional impairments, we isolated hindlimb extensor digitorum longus (EDL) muscles from P56 mice and conducted electrophysiological recording. We incubated the muscles with μ-conotoxin, which selectively inhibits muscle voltage-gated Na+ channels, prevented the induction of muscle action potential (Ling et al., 2010; Zanetti 2018), and recorded the Miniature endplate potential (mEPP) and evoked endplate potential (eEPP). (Ling et al., 2010; Zanetti 2018). mEPP, a response that occurs when spontaneously released acetylcholine binds to nicotinic AChR without nerve stimulation, was measured ex vivo near the NMJs of the EDL muscles in the control and SMN2 1-copy Smn^ΔMPC^ mice (Figure 5A). The mEPP amplitude was increased in SMN2 1-copy Smn^ΔMPC^ mice (Figure 5B), whereas mEPP frequency was comparable between the controls and mutants (Figure 5C). The results indicate that the NMJ synapses of SMN2 1-copy Smn^ΔMPC^ mice are functional and more sensitive to acetylcholine compared to the controls. Next, we measured the eEPP by stimulating an action potential at the peroneal nerve (Figure 5D). Despite the increased mEPP amplitude, the amplitude of eEPPs was significantly decreased in SMN2 1-copy Smn^ΔMPC^ mice (Figure 5E). These results suggest that the nerve terminals in SMN2 1-copy Smn^ΔMPC^ mice exhibit decreased quantal content. This could be due to a decrease in vesicle release probability. Notably, there was no difference in the paired-pulse response, indicating normal neurotransmitter release probability (Figure 5F-G). Taken together, these findings suggest that the presynaptic neurotransmission ability of the NMJ is reduced in SMN2 1-copy Smn^ΔMPC^ mutants.

**Figure 5.**
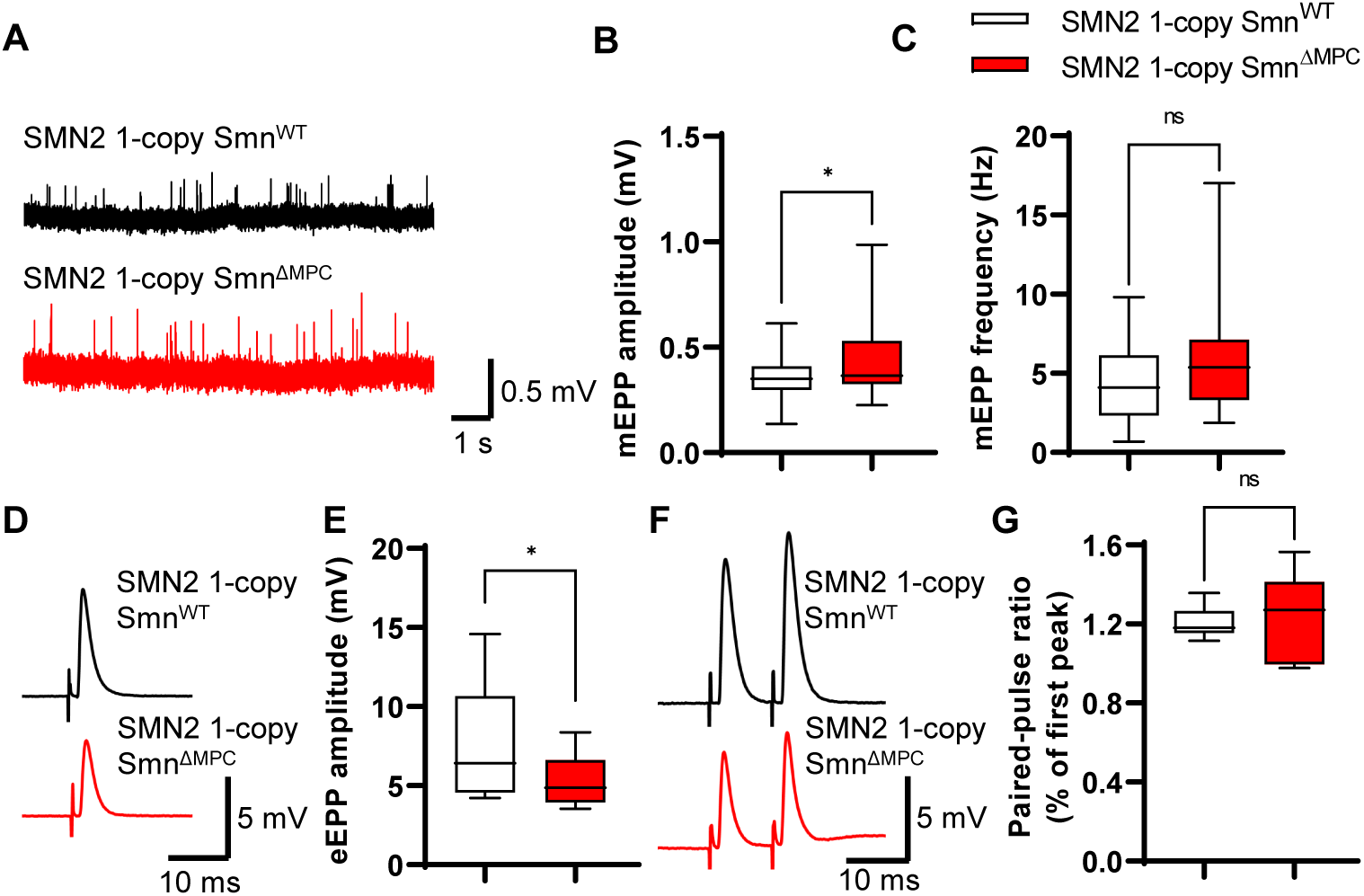
Reduced presynaptic neurotransmission ability in the NMJs of SMN2 1-copy Smn^ΔMPC^ mice. **(A)** Representative traces of mEPP from SMN2 1-copy Smn^ΔWT^ (top) and SMN2 1-copy Smn^ΔMPC^ (bottom) mice. **(B-C)** SMN2 1-copy mutant’s NMJs showed an increase in mEPP amplitude and no differences in mEPP frequency (1-copy control, n = 25, 9 mice; 1-copy mutant, n = 21, 8 mice; Unpaired t-test with Welch’s correction). **(D)** Representative traces of eEPP from SMN2 1-copy Smn^ΔWT^ (top) and SMN2 1-copy Smn^ΔMPC^ (bottom) mice. **(E)** The mutant’s NMJs showed a stronger amplitude of eEPPs (1-copy control, n = 12, 4 mice; 1-copy mutant, n = 12, 3 mice; Unpaired t-test with Welch’s correction). (**F**) Representative traces of paired-pulse response from SMN2 1-copy Smn^ΔWT^ (top) and SMN2 1-copy Smn^ΔMPC^ (bottom) mice. **(G)** Paired-pulse response was not different between SMN2 1-copy control and mutant NMJs, indicating a comparable neurotransmitter release probability (1-copy control, n = 8, 3 mice; 1-copy mutant, n = 6, 3 mice; Unpaired t-test with Welch’s correction). The electrophysiological recording was performed in the EDL muscle at P56. ns; not significantly different. \**p* < 0.05. All box-and-whisker plots show the median, interquartile range, minimum, and maximum. For the box-and-whisker plots, range bars show minimum and maximum (**B, C, E, G**).

### Disturbed nerve terminal structure in SMN2 1-copy Smn**^Δ^**^MPC^ mice

To examine the NMJ ultrastructure of SMN2 1-copy Smn^ΔMPC^ mutants, we utilized transmission electron microscopy (TEM) (Figure 6A). The density of junctional folds in SMN2 1-copy Smn^ΔMPC^ mutant specimens was comparable to that of the control (Figure 6B). However, the density of synaptic vesicles was substantially elevated in the SMN2 1-copy mutants (Figure 6C). Since previous electrophysiological results suggested a decrease in presynaptic neurotransmission capacity in SMN2 1-copy mutants, this could be due to synaptic vesicles failing to fuse with the membrane, leading to the accumulation of vesicles in the terminal and reduced quantal contents in Figure 5. In Figure 3H, larger synaptophysin coverage in the mutant may be caused by this synaptic vesicle accumulation. Additionally, the detachment of the nerve terminal is more frequent at the NMJ of mutants (Figure 6D). The detachment of nerve terminals observed in SMN2 1-copy Smn^ΔMPC^ mutants could have also resulted in diminished presynaptic neurotransmission capacity. Collectively, these findings indicate that SMN2 1-copy Smn^ΔMPC^ mutants have nerve terminal-specific pathological defects at the NMJ ultrastructural level.

**Figure 6.**
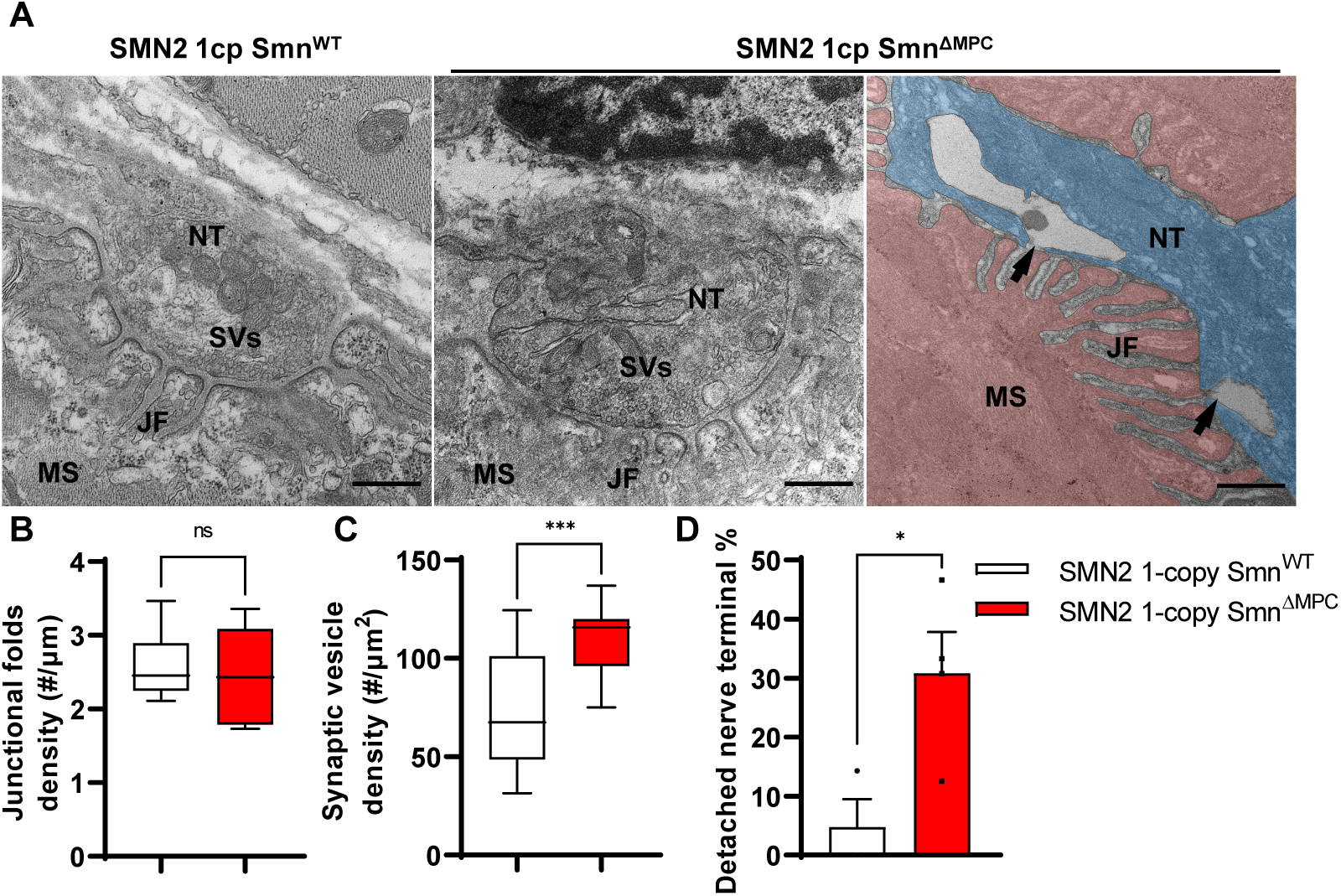
Abnormal nerve terminal ultrastructure in SMN2 1-copy mutant. **(A)** Representative TEM images from NMJs of SMN2 1-copy Smn^ΔWT^ and SMN2 1-copy Smn^ΔMPC^ mice at P56. Scale bars, 500 nm. Nerve terminal (NT; indicated by the blue zone). Synaptic vesicles (SVs). Muscle fiber (MS; indicated by the red zone). Endplate junctional folds (JF). Nerve terminal detachment (indicated by arrow) was observed in SMN2 1-copy Smn^ΔMPC^ mice. **(B)** The density of junctional folds in the NMJ of SMN2 1-copy Smn^ΔMPC^ mice no significant change compared to the control, whereas (**C**) the density of synaptic vesicles was increased (n = 3-4 mice in each genotype; Unpaired t-test with Welch’s correction). (**D**) The detachment of the nerve terminal occurs more frequently at the NMJ of mutants (n = 3-4 mice in each genotype; Unpaired t-test with Welch’s correction). ns; not significantly different. \**p* < 0.05; \*\*\**p* < 0.001. All box-and-whisker plots show the median, interquartile range, minimum, and maximum. For the box-and-whisker plots, range bars show minimum and maximum (**B, C**). For the bar graph, error bars show s.e.m(**D**).

### FAPs transplantation rescues NMJ morphology in limb mesenchymal SMN mutants

SMN-deleted limb mesenchymal tissues in SMN2 1-copy Smn^ΔMPC^ mutants comprise not only FAPs, but also bone, cartilage, pericytes, and tendon, among others (Leinroth et al., 2022; Nassari et al., 2017). To evaluate the critical role of FAPs in the postnatal development of the NMJ, we isolated fluorescent protein-labeled wild-type FAPs from *Prrx1^Cre^; Rosa26^YFP/+^*or *Prrx1^Cre^; Rosa26^tdTomato/+^*mice and transplanted them into the TA muscles of SMN2 1-copy Smn^ΔMPC^ mice at postnatal day 10. The TA muscle on the contralateral side was treated with a vehicle as a control. In SMN2 1-copy Smn^ΔMPC^ mice at P56, the muscle that received FAPs showed decreased presynaptic fragmentation, NF varicosities, and postsynaptic fragmentation compared to the contralateral muscle (Figure 7A-D). As a result, the transplanted muscle was not significantly different from the control except for NF varicosities. These data demonstrate that the transplantation of wild-type FAPs rescues the abnormal NMJ development in SMN2 1-copy Smn^ΔMPC^ mice. Overall, our findings indicate that SMN depletion in FAPs leads to the neuronal SMN-independent NMJ pathology in severe SMA, which is rescued by the transplantation of healthy FAPs.

**Figure 7.**
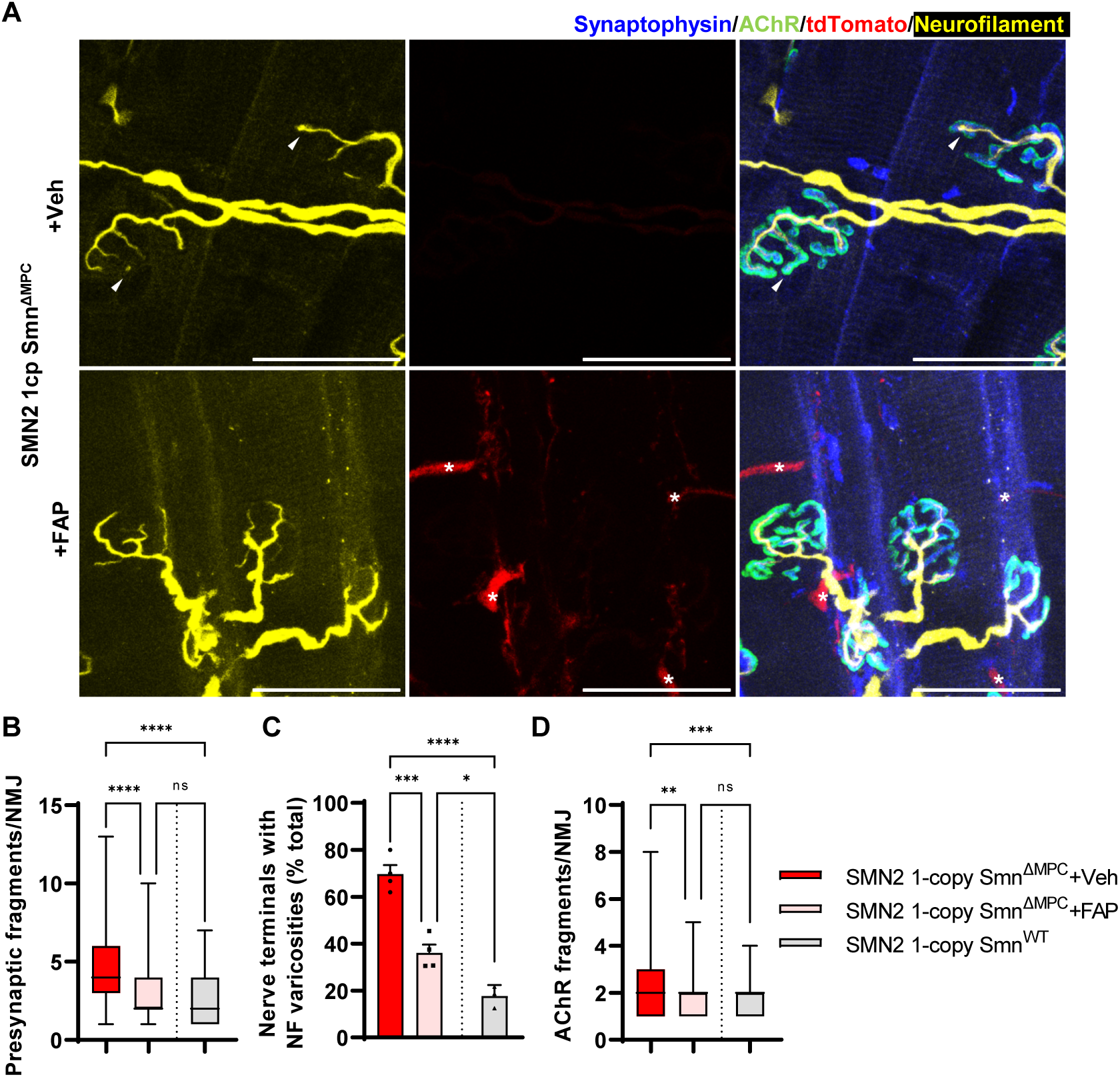
Improved postnatal NMJ development in the TA muscle of SMN2 1-copy Smn^ΔMPC^ mice following healthy FAPs transplantation. (**A**) Immunostaining of NMJs in tdTomato^+^ FAPs-transplanted TA muscle (+FAP) and vehicle-treated contralateral muscle (+Veh) in SMN2 1-copy Smn^ΔMPC^ mice at P56 with anti-NF (Yellow), anti-synaptophysin (Blue), α-Btx staining AChR (Green), and tdTomato fluorescence (Red). Scale bars, 40 μm. The images revealed NF varicosities (indicated by arrowheads) in the +Veh NMJs. The tdTomato+ FAPs (marked by asterisks) were transplanted into +FAP NMJs, which exhibited (**B**) decreased presynaptic fragmentation, (**C**) NF varicosities, and (**D**) AChR fragmentation compared to +Veh NMJs and similar to wild-type NMJs. (n = 3-4 mice in each group; Unpaired t-test with Welch’s correction). ns; not significantly different. \*\**p* < 0.01, \*\*\**p* < 0.001, \*\*\*\**p* < 0.0001. All box-and-whisker plots show the median, interquartile range, minimum, and maximum. For the box-and-whisker plots, range bars show minimum and maximum (**B, D**). For the bar graph, error bars show s.e.m (C).

## Discussion

In this paper, we elucidate the contribution of SMN depletion in mesenchymal progenitors for the pathogenesis of SMA. To test this hypothesis, we generated conditional knockout mouse strains to delete the Smn allele specifically in limb mesenchymal cells and carry human *SMN2* copies. Our research using these mouse models resulted in three major discoveries. First, SMN deficiency in FAPs contributes to NMJ pathological defects in SMA. We observed delayed NMJ maturation and varicosities in juvenile SMN2 1-copy Smn^ΔMPC^ mutant. The pathogenic NMJ phenotypes were also observed in the SMAΔ7 mutant, which is one of the severe SMA mouse models (Kong et al., 2009; Martinez, 2012; Kariya et al., 2008). As the SMAΔ7 mutant typically lives for approximately 12 days, the fragmentation of the NMJ in adult SMN2 1-copy mutant was not evaluated in the severe SMA mutant. Nevertheless, models that induce a conditional adult SMN deficiency through either Cre^ER^ allele or oligonucleotide administration resulting in SMN reduction, demonstrated the depletion of SMN throughout the body caused fragmentation of NMJ (Sahashi et al., 2013; Kariya et al., 2014). Thus, we demonstrate that the SMN2 1-copy Smn^ΔMPC^ mutant model mimics whole-body SMA mouse models in NMJ morphology. In the electrophysiological test, the SMAΔ7 mutant exhibited reduced quantal content, readily releasable pool, and vesicle release probability (Torres-Benito et al., 2011). In our study, we did not directly assess the readily releasable pool in SMN2 1-copy Smn^ΔMPC^ mutant by stimulus train of electrophysiological recording but instead showed reduced quantal content and normal vesicle release probability. In previous studies, it was theorized that the decrease in quantal contents in SMAΔ7 mutants resulted from decreased synaptic vesicle density at nerve terminals caused by motor neuron defects and abnormal axonal transport (Dale et al., 2011; Kong et al., 2009). However, we found that synaptic vesicle density was increased in the SMN2 1-copy Smn^ΔMPC^ mutant with SMN-sufficient motor neurons. It is possible that the alteration of active zones, which were also altered in the SMA motor terminals (Kong et al., 2009), contributed to the reduction in synaptic vesicle fusion and the decreased quantal contents. Indeed, nerve terminal detachment in SMN2 1-copy mutant mice was also found in active zone complex protein integrin-α3 knockout mice (Ross et al., 2017). Based on the rescue data of transplanted healthy FAPs, we can report that FAPs-specific SMN depletion is involved in NMJ pathology of SMA.

Second, we demonstrated that skeletal growth defects, a phenotype observed in SMA (Khatri et al., 2008; Vai et al., 2015; Wasserman et al., 2017; Baranello et al., 2019; Hensel et al., 2020), are a cell-autonomous pathological effect of the depletion of SMN in chondrocytes. We observed reduced bone size and volume in juvenile SMN2 1-copy Smn^ΔMPC^ mutant. Depletion of SMN from chondrocytes is sufficient to cause growth problems with chondrocyte-secreted IGF defects, which affects growth plate homeostasis. Previous studies have reported that *IGF1* overexpression improves biochemical and behavioral manifestations in SMA mice, suggesting potential therapies for SMA (Bosch-Marce et al., 2011; Tsai et al., 2014). Our study showed that these IGF therapies for SMA could be one to consider for treating bone growth abnormalities. In addition, the skeletal growth defect may affect physiologic muscle atrophy through imbalanced muscle contraction and reduced tension. This physiologic atrophy may be part of SMA-associated muscle weakness independent of neurogenic atrophy.

Third, it was demonstrated that adequate levels of SMN protein are essential for MPCs to contribute to limb neuromusculoskeletal development. The mutants with only 1 copy of *SMN2* exhibit problematic symptoms observed in both SMA patients and mouse models, while SMN2 2-copy mutants display a typical phenotype in bone and NMJ. The lack of SMN protein in MPCs by insufficient *SMN2* copies, similar to the deficiency seen in severe SMA, is responsible for the onset of SMA pathology. Thus, we propose that restoration of deficient SMN in MPCs is crucial for rehabilitating their function. Based on these discoveries, SMN replenishment treatments for MPCs, specifically FAPs, and chondrocytes, are necessary to provide a complete solution for neuromusculoskeletal defects in severe SMA patients.

The initial focus for the treatment of SMA was on motor neurons located in the spinal cord, and pharmaceuticals were created to address the deficiency of SMN in these neurons (Mercuri et al., 2020). For example, intrathecal injections of Spinraza can efficiently boost SMN in the CNS, including the spinal cord (Passini et al., 2011; Claborn et al., 2018). However, Spinraza does not address the lack of SMN in peripheral tissues, including mesenchymal cells, highlighted in recent studies. The drug Zolgensma, which employs AAV9 to express SMN through systemic delivery, appears to resolve this issue (Foust et al., 2009; Foust et al., 2022; Valori et al., 2010; Mattar et al., 2013). Nevertheless, a prior study indicates that chondrocytes within the growth plate and articular cartilage do not get infected with AAV9 (Yang et al., 2019). Furthermore, it is well-established that DNA vectors delivered via AAV9 undergo dilution during cell proliferation (Penaud-Budloo et al., 2008; Colella et al., 2018; Van-Alstyne et al., 2021; Heller et al., 2021). Given the infection issue and the dilution issue by the active cell population changes of chondrocytes and FAPs during postnatal development (Petrany., 2020; Bachman et al., 2022), SMN supplementation through Zolgensma alone would not be sufficient for severe SMA patients. Thus, there is an urgent need to research and develop therapeutic strategies that target mesenchymal progenitors.

While this study is the first to demonstrate the impact of SMN depletion in FAPs on the NMJ development, it does not elucidate the specific mechanism by which FAPs influence the NMJ. Our study observed NMJ recovery in the muscles transplanted with healthy FAPs, but not in the contralateral muscles, indicating that FAPs are likely involved in NMJ development through juxtacrine or paracrine signaling. Since FAPs interact with surrounding tissues through a variety of signaling factors, such as extracellular matrix (Contreras et al., 2021; Scott et al 2019), Wnt-related protein (Lukjanenko et al., 2019), and Bmp signaling protein (Uezumi et al., 2021; Camps et al., 2020), mis-splicing of the signaling factors due to SMN reduction could disturb the homeostasis of neighboring tissues (Zhang et al., 2008). Furthermore, the *Hsd11b1*-positive subpopulation of FAPs associated with NMJ was discovered through single-cell RNA sequencing in a recent study (Leinroth et al., 2022). This population is located adjacent to the NMJ and responds to denervation, indicating an increased possibility of interaction with the NMJ organization. Therefore, it is necessary to conduct additional investigations into the expression of various signaling factors by diverse FAP subpopulations in future studies.

## Methods

### Animals

*Prrx1^Cre^* (stock 005584), *Smn^f7/+^* (stock 006138), *Rosa26^YFP/+^* (stock 006148), *Rosa26^tdTomato/+^* (stock 007914), *SMN2*^+/+^, *Smn*^+/-^ (stock 005024 – *Smn* knockout and *SMN2* homologous transgenic mouse –) and *Smn^+/-^; SMN2^+/+^; SMN*Δ*7^+/+^* (stock 005025 005024 – *Smn* knockout and *SMN2*, *SMN*Δ*7* homologous transgenic mouse –) mice were acquired from The Jackson Laboratory (Bar Harbor, ME, USA). SMN2 0-copy Smn^ΔMPC^ mice (*Prrx1^Cre^; Smn^f7/f7^*), SMN2 copy Smn^ΔMPC^ mice (*Prrx1^Cre^; Smn^f7/f7^; SMN2^+/0^* –*SMN2* heterologous allele–), and SMN2 copy Smn^ΔMPC^ mice (*Prrx1^Cre^; Smn^f7/f7^; SMN2^+/+^*–*SMN2* homologous allele–),) were generated by crossing *Prrx1^Cre^* mice with *Smn^f7/+^* and *Smn^+/-^; SMN2^+/+^*mice. To utilize fibro-adipogenic progenitors (FAPs) transplantation, we generated *Prrx1^Cre^; Rosa26^YFP/+^* mice and *Prrx1^Cre^; Rosa26^tdTomato/+^* mice by breeding *Prrx1^Cre^* with *Rosa26^YFP/+^* and *Rosa26^tdTomato/+^*mice, respectively. To avoid deletion of the floxed allele by *Prrx1^Cre^*expression in female germline, we bred females without *Prrx1^Cre^*line to *Prrx1^Cre^* transgenic males. SMAΔ7 mutants (*Smn^-/-^; SMN2^+/+^; SMN*Δ*7^+/+^*) were produced by mating *Smn^+/-^; SMN2^+/+^; SMN*Δ*7^+/+^* mice. Both male and female mice were used in the experiments, and no sex-specific differences were observed. Control littermates lacking *Prrx1^Cre^* were utilized for analysis. All mouse lines were housed under controlled conditions with specific pathogen-free and handled according to the guidelines of the Seoul National University Institutional Animal Care and Use Committee (IACUC).

### Micro CT

The femurs from three groups - control, SMN2 2-copy Smn^ΔMPC^, and SMN2 1-copy Smn^ΔMPC^ mice at postnatal day 14 (P14) - were isolated and cleaned of muscles and skin. Subsequently, the femurs were preserved in 4% paraformaldehyde (PFA) in PBS overnight at 4°C before micro-computed tomography (micro CT). The femurs were then imaged through a Quantum GX II micro-CT imaging system (PerkinElmer, Hopkinton, MA, USA). The X-ray source for scanning was set at 90 kV and 88 mA with a field of view of 10 mm (voxel size, 20 μm; scanning time, 14 min). The 3D imaging was viewed using the 3D Viewer software of the Quantum GX II. The size and volume of the femur bone were measured via AccuCT analysis software within the ossified diaphysis and metaphysis of the femur, excluding the epiphysis. Bone mineral density (BMD) calibration was performed using a 4.5 mm BMD phantom and BMD measurements were taken at the center of the diaphysis.

### Histology

Alcian blue & alizarin red staining was performed on control and SMN2 0-copy Smn^ΔMPC^ mice at E18.5 using a previously reported protocol (Ovchinnikov, 2009). For hematoxylin & eosin and toluidine blue staining of the growth plate in the control and SMN2 1-copy Smn^ΔMPC^ mice at P14, the femurs were fixed in 4% paraformaldehyde for 24 hours, rinsed in running tap water for 24 hours, and incubated with 10% EDTA (pH 7.4) at 4°C with shaking for 2-3 days. Subsequently, the samples were rinsed in running tap water for 24 hours and then dehydrated through ethanol/xylene and embedded in paraffin. The embedded samples were then sectioned to a thickness of 5 µm, rehydrated, and stained with Gill No. 3 formula hematoxylin and eosin Y (H&E, Sigma Aldrich, St. Louis, MO, USA) and toluidine blue (Sigma Aldrich, St. Louis, MO, USA). Stained slides were analyzed using 10× and 20x objectives in the EVOS M7000 imaging system (Thermo Fisher Scientific, Waltham, MA, USA). The growth plate’s proliferative and hypertrophic zones were defined by their respective cell sizes.

### Hindlimb fibro-adipogenic progenitors isolation

Isolation of limb muscle FAPs was performed according to a previously reported protocol (Kim et al., 2021) with modifications. Limb muscles were dissected and mechanically dissociated in Dulbecco’s modified Eagle’s medium (DMEM, Hyclone) containing 10% horse serum (Hyclone, Logan, UT, USA), collagenase II (800 units/mL; Worthington, Lakewood, NJ, USA), and dispase (1.1 units/mL; ThermoFisher Scientific, Waltham, MA, USA) at 37°C for 40 min. Digested suspensions were subsequently triturated by sterilized syringes with 20G 1/2 needle (BD Biosciences, Franklin Lakes, NJ, USA) and washed with DMEM to harvest mononuclear cells. Mononuclear cells were stained with corresponding antibodies. All antibodies used in FACS analysis were listed in Supplemental Table 1. To exclude dead cells, 7AAD (Sigma Aldrich; St. Louis, MO, USA) was used. Stained cells were analyzed and 7AAD^−^Lin^−^Vcam^−^Sca1^+^ (Stem cell antigen 1; *Ly6a*) (FAPs) were isolated using FACS Aria III cell sorter (BD Biosciences) with 4-way-purity precision. For western blot, freshly isolated FAPs were cultured at 37 °C in alpha-MEM (Hyclone) supplemented with Antibiotic-Antimycotic (anti-anti, Gibco) and 20% FBS (Hyclone). For transplantation, YFP^+^ FAPs and tdTomato^+^ FAPs were isolated from postnatal day 10∼21 *Prrx1^Cre^; Rosa26^YFP/+^* mice and *Prrx1^Cre^; Rosa26^tdTomato/+^* mice, respectively. Isotype control density plots were used as a reference for positive gating.

### Chondrocytes isolation

Isolation of chondrocytes followed a modified protocol previously published (Jonason et al., 2015). Using blunt forceps, cartilage caps were removed from P21 mouse femoral heads and dissected into ∼1mm fragments in a petri dish with 10X anti-anti in PBS. The cartilage fragments were washed twice with PBS and then incubated in 5 mL collagenase II solution (800 units/mL collagenase II in DMEM with anti-anti, sterilized by 0.2 µm filtration) in a 60-mm culture dish at 37 °C in a 5% CO2 incubator overnight. Chondrocytes were released by pipetting the remaining cartilage fragments ten times and then filtering them through a 70 μm cell strainer to a 50 ml conical tube. The cells were then washed twice with PBS and pelleted by centrifugation at 500g for 5 min. Subsequently, the cells were cultured overnight at 37 °C in a 5% CO2 incubator, in complete culture medium (DMEM with 10% FBS and anti-anti) overnight at 37 °C in a 5% CO2 incubator.

### PCR reaction

To detect the presence of Smn exon 7 floxed and deleted allele, 20 mg of each tissue were dissolved in direct PCR buffer (VIAGEN) with proteinase K overnight at 65°C. After inactivation at 95°C for 30 minutes, PCR was performed using previously reported primer sets to verify the existence of Cre, Smn^f7^, and Smn^Δ7^ alleles (Frugier et al., 2000). Primers are listed in Supplemental Table 1.

### RNA extraction and measurement of mRNA expression

Total RNA was extracted from the brain, liver, spinal cord L4, tibialis anterior muscle, freshly isolated FAPs, and chondrocytes using a TRIzol Reagent (Life Technologies, Carlsbad, CA, USA) and analyzed by qRT-PCR. First-strand complementary DNA was synthesized from 1 μg of RNA using ReverTra Ace (Toyobo, Osaka, Japan) containing random oligomer according to the manufacturer’s instructions. qRT-PCR (Qiagen) was performed with SYBR Green technology (SYBR Premix Ex Taq, Qiagen) using specific primers against indicated genes. Relative mRNA levels were determined using the 2^-ΔΔCt^ method and normalized to *Gapdh* (Figure 1H, Figure 1—figure supplement 1K). Primers are listed in Supplemental Table 1.

### Western blot

Cultured FAPs at passage 3 were homogenized in RIPA buffer (50 mM Tris-HCl, pH 7.5, 0.5% SDS, 20 µg/ mL aprotinin, 20 µg/ mL leupeptin, 10 µg/ mL phenylmethylsulfonyl fluoride, 1 mM sodium orthovanadate, 10 mM sodium pyrophosphate, 10 mM sodium fluoride, and 1 mM dithiothreitol). Cell lysates were centrifuged at 13,000 rpm for 15 min. Supernatants were collected and subjected to immunoblot. BCA protein assay (Thermo Fisher Scientific) was used for estimating total protein concentrations. Normalized total proteins were analyzed by electrophoresis in 10% polyacrylamide gels and transferred to PVDF membranes (Millipore, Billerica, MA, USA). Membranes were blocked in 5% skim milk (BD Biosciences) in TBS with 0.1% Tween-20 and incubated with primary antibodies overnight at 4°C. After incubation with the corresponding HRP-conjugated secondary antibodies, the membranes were developed using a Fusion solo chemiluminescence imaging system (Vilber, Marne-la-Vallée, France). α-tubulin was used as a loading control. Band intensities were quantified using ImageJ software. Antibodies used in this study were listed in Supplemental Table 1. Primary and secondary antibodies were diluted 1:1000 and 1:10000 with PBS containing 0.1% Tween-20 and 3% BSA, respectively.

### FAPs transplantation

FAPs transplantation was performed according to a previously reported protocol (Kim et al., 2021) with modifications. YFP^+^ or tdTomato^+^ FAPs (7AAD^−^Lin^−^Vcam^−^Sca1^+^) were isolated by FACS from the limb muscles of indicated mice. 1×10^5^ FAPs were suspended in 0.1% gelatin (Sigma Aldrich; St. Louis, MO, USA) in PBS and then transplanted into one side of the TA muscles of SMN2 1-copy Smn^ΔMPC^ mice. The contralateral muscle received an equivalent volume of 0.1% gelatin in PBS (Vehicle).

### Electrophysiology

The extensor digitorum longus (EDL) muscle was dissected from control and SMN2 1-copy Smn^ΔMPC^ mice, along with the peroneal nerve, and then pinned to a Sylgard-coated recording chamber. Intracellular recording was conducted in oxygenated Ringer’s solution, which comprised 138.8 mM NaCl, 4 mM KCl, 12 mM NaHCO3, 1 mM KH2PO4, 1 mM MgCl2 and 2 mM CaCl2 with a pH of 7.4. Action potential of the muscle was prevented by preparing the muscle in 2.5 uM μ-conotoxin GIIB (Alomone, Jerusalem, Israel) for 10 minutes beforehand. The recording was performed in toxin-free Ringer’s solution. Miniature endplate potentials (mEPPs) were recorded from a junction, followed by recordings of evoked endplate potentials (eEPPs) by stimulating the attached peroneal nerve. The eEPPs were elicited using evoked stimulation. Paired-pulse stimulation (10 ms inter-stimulus interval) was utilized to assess synaptic transmission. The data was obtained and analyzed with Axoclamp 900A and Clampfit version 10.7 software.

### Transmission electron microscopy

NMJ transmission electron microscopy followed a modified protocol previously reported (Modla et al., 2010). Mouse EDL muscle was swiftly excised and fixed in 4% paraformaldehyde (PFA) dissolved in Sorensen’s phosphate buffer (0.1M, pH 7.2), followed by washing in 0.1 M phosphate buffer. The EDL was then gradually infiltrated on a rotator at room temperature with sucrose: 0.1M phosphate buffer solutions of 30% and 50%, for one hour each, followed by an overnight incubation in 70% sucrose. Excess sucrose was then eliminated using filter paper, and the muscle was embedded in an optimal cutting temperature compound (O.C.T.; Sakura Fineteck, Torrance, CA, USA), followed by being frozen in a cryostat (Leica, Wetzlar, Germany). 10-µm thick longitudinal sections were washed in PBS and treated with Alexa fluor 555-conjugated α-bungarotoxin (1:500, Invitrogen) for an hour. Imaging was conducted with the EVOS M7000 imaging system, and we selected 4-5 NMJ-rich regions for processing with TEM. The sections were fixed with 2% glutaraldehyde and 2% PFA in 0.1M cacodylate buffer (pH 7.2) for two hours at room temperature, with an additional overnight incubation at 4L. After washing with 0.1M cacodylate buffer they were post-fixed with 1% osmium tetraoxide in 0.1M cacodylate buffer (pH 7.2)-for two hours at 4°C. The sections were then stained en bloc with 0.5% uranyl acetate overnight, washed with distilled water, and dehydrated using serial ethanol and propylene oxide. The sections were embedded in epoxy resin (Embed-812, Electron Microscopy Sciences) and detached from the slides by dipping them in liquid nitrogen. Ultra-thin sections (70nm) were prepared with a diamond knife on an ultramicrotome (ULTRACUT UC7, Leica) and mounted on 100 mesh copper grids. Sections were stained with 2% uranyl acetate for 10 minutes and lead citrate for 3 minutes, then observed using a transmission electron microscope (80kV, JEM1010, JEOL or 120kV, Talos L120C, FEI). Synaptic vesicle density was quantified within a distance of 500 nm from the presynaptic membrane.

### Immunohistochemistry

For NMJ staining, freshly dissected TA muscles were fixed in 4% PFA for 30 min at room temperature. Subsequently, the muscles were cryoprotected in 30% sucrose overnight, embedded in O.C.T., snap-frozen in liquid nitrogen, and stored at -80°C prior to sectioning. Longitudinal 40 μm-thick sections were obtained from the embedded muscles using a cryostat. The sections were blocked for 2 hours at room temperature using 5% goat serum and 5% bovine serum albumin in PBS/0.4% Triton X-100. Then, the sections were incubated with primary antibodies in the blocking buffer for two days at 4 °C. After washing the sections three times with PBS/0.4% Triton X-100, the sections were stained with secondary antibodies overnight at 4 °C, and then incubated with Alexa fluor 488 or 555-conjugated α-bungarotoxin (1:500, Invitrogen) for 2hr at RT, washed three times with PBS/0.4% Triton X-100 and mounted in VECTASHIELD. Z-serial images were collected at 40X with a Leica SP8 confocal laser scanning microscope. To analyze NMJ morphology, LasX software was used to obtain maximal projections. Neurofilament (NF) varicosity refers to the varicose NF end connected to the rest of the nerve terminal. To quantify NMJ size and synaptophysin coverage, the Btx area and synaptophysin area were measured by ImageJ analysis software.

For bone section staining, the 5 µm-thick bone sections were rehydrated and antigen retrieval was then performed in citrate buffer (10 mM citric acid, pH 6) at 95°C 20 min. The sections were blocked for 1 hr at room temperature using 5% goat serum and 5% bovine serum albumin in PBS/0.4% Triton X-100. Then, the sections were incubated with primary antibodies in the blocking buffer at 4 °C overnight. After washing the sections three times with PBS/0.1% Triton X-100, the sections were stained with secondary antibodies for 1 hr at RT, washed and mounted. Imaging was conducted with the EVOS M7000 imaging system.

### Statistical analysis

All statistical analyses were performed using GraphPad Prism 9 (GraphPad Software). For comparison of significant differences in multiple groups for normally distributed data, statistical analysis was performed by one-way or two-way ANOVA followed by Tukey’s pairwise comparison post hoc test. For non-normally distributed data, Brown–Forsythe and Welch ANOVA followed by Games-Howell multiple comparisons test was used. For the comparison of two groups, Student’s unpaired t-test assuming a two-tailed distribution with Welch’s correction was used. Unless otherwise noted, all error bars represent s.e.m. The number of biological replicates and statistical analyses for each experiment were indicated in the figure legends. Independent experiments were performed at least in triplicates. P < 0.05 was considered statistically significant at the 95% confidence level. \**p* < 0.05, \*\**p* < 0.01, \*\*\**p* < 0.001, \*\*\*\**p* < 0.0001.

## Supplemental information

Supplemental information includes one Supplementary Table.

## Ethics

The care and treatment of animals were approved by the IACUC of Seoul National University.

## Acknowledgments

We express our gratitude to the Kong and Choi laboratory members for their valuable feedback during the project. This work was supported by the National Research Foundation of Korea (NRF-2022R1A2C3007621 and NRF-2020R1A5A1018081).

---

**Figure 1 -figure supplement 1.**
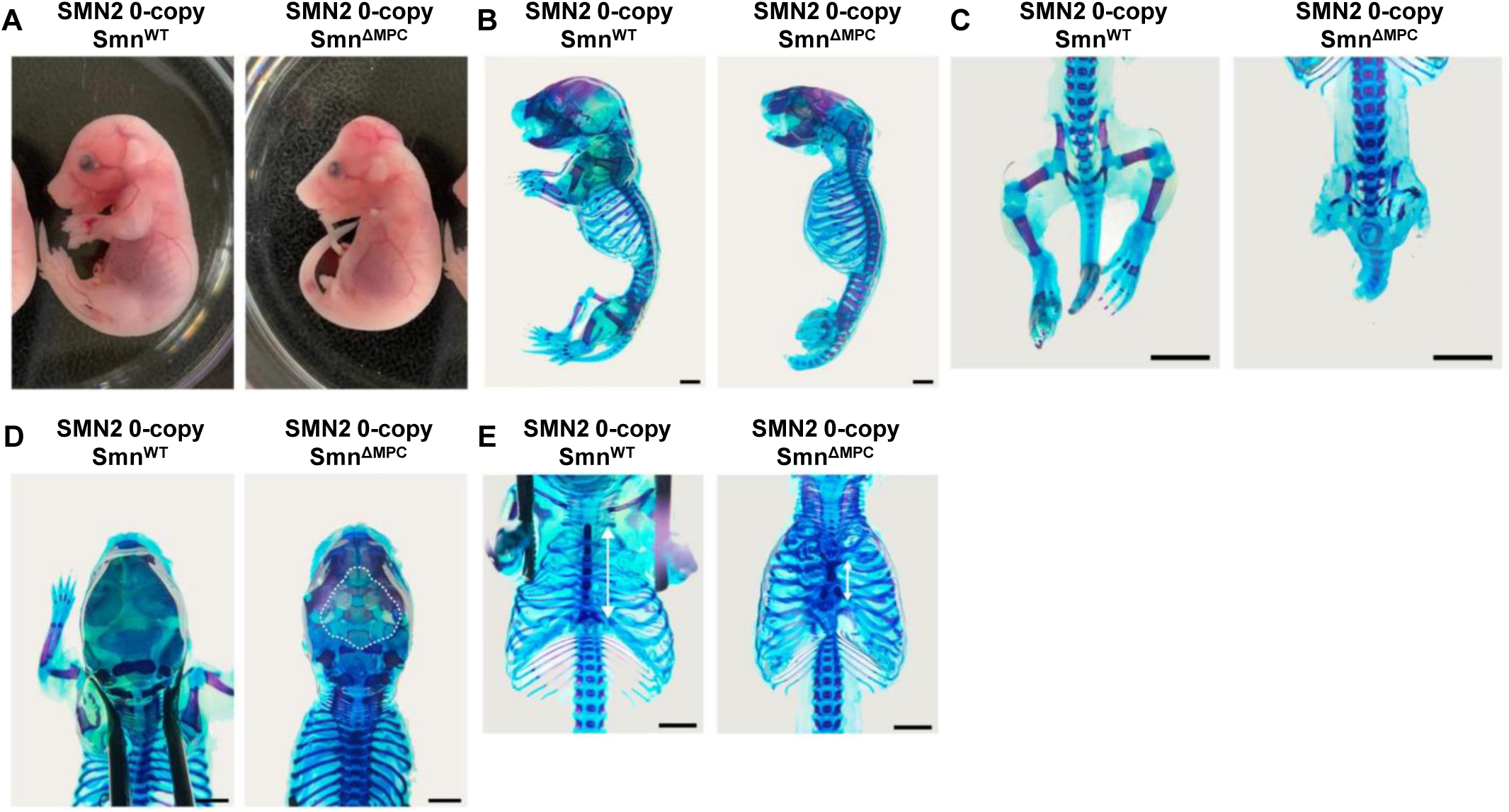
Growth defects in the Prrx1-lineage bone of SMN2 0-copy Smn^ΔMPC^ mice. (**A)** Representative control and mutant mice with 0 copies of SMN2 were photographed at E18.5. (**B-C**) Alcian blue & alizarine red staining in the SMN2 0-copy Smn^ΔMPC^ mutant showed abnormal skeletal development of the limbs, (**D**) calvaria (shown by the dashed line), and (**E**) sternum (indicated by a white arrow). Scale bars, 5 mm (**B-C**) and 2 mm (**D-E**).

**Figure 1 -figure supplement 2.**
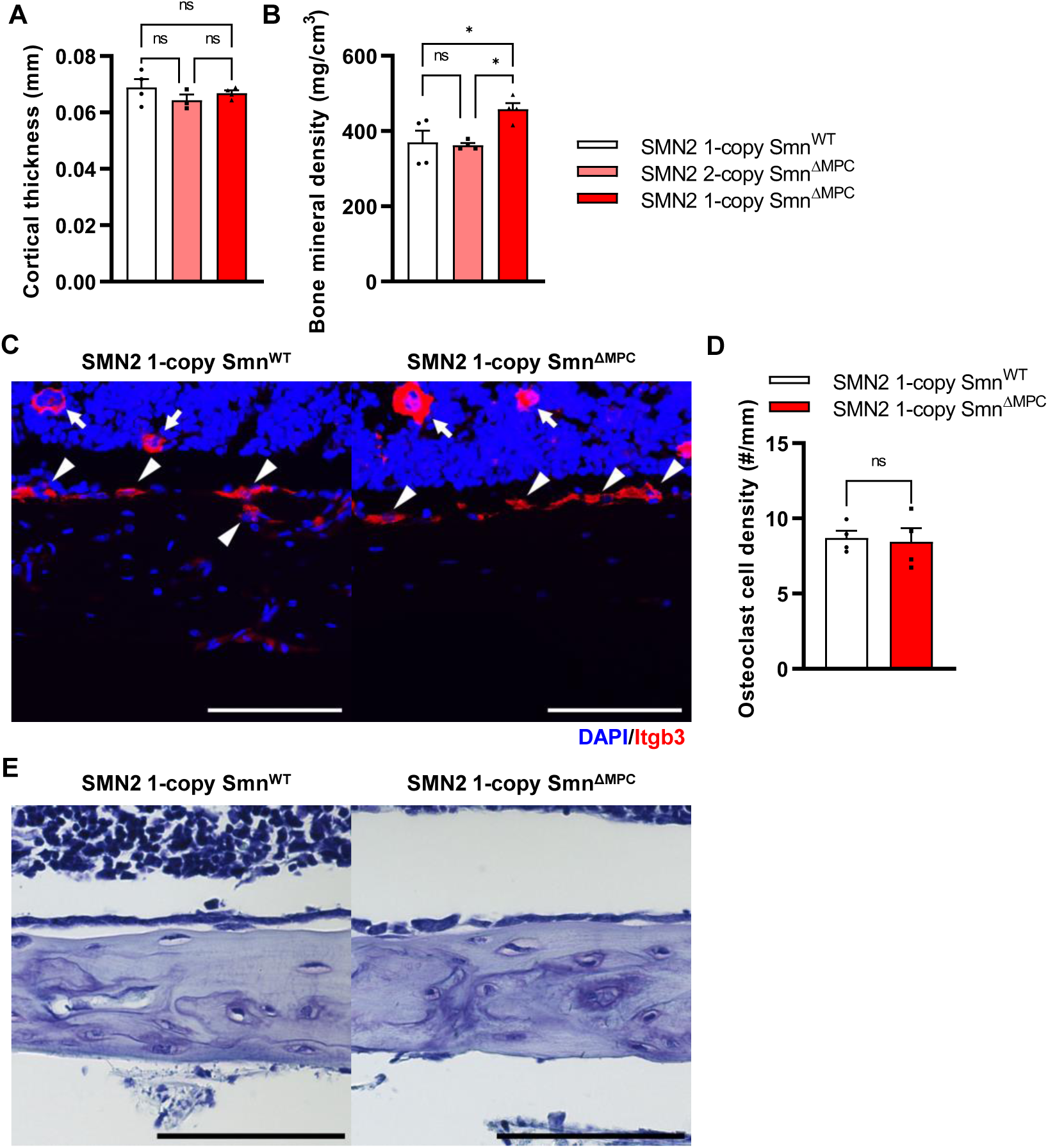
Osteoclasts and osteoblasts were undisturbed in SMN2 1-copy Smn^ΔMPC^ mice. (**A**) Cortical bone thicknesses were not significantly different between the control and mutant groups. (**B**) Bone mineral density was slightly increased in the diaphysis of SMN2 1-copy Smn^ΔMPC^ mice. The micro CT analysis was performed in femur diaphysis and metaphysis from SMN2 1-copy Smn^WT^, SMN2 2-copy, and 1-copy Smn^ΔMPC^ mice at P14. 1-way ANOVA with Tukey’s post hoc test, n = 3-5 mice in each genotype (**A-B**). (**C**) Representative images of osteoclast marker *Itgb3* immunostaining in femur diaphysis cortical bone from mice at P14. Scale bars, 100 μm. Itgb3+ hematopoietic cell (indicated by arrow) and osteoclast (indicated by arrowhead) were imaged. (**D**) It showed similar osteoclast density. n = 3 mice in each genotype; Unpaired t-test with Welch’s correction. (**E**) Representative images of toluidine blue staining in femur diaphysis cortical bone for osteoblast evaluation. Scale bars, 100 μm. ns; not significantly different. \**p* < 0.05. Error bars show s.e.m.

